# The *shh* limb enhancer is activated in patterned limb regeneration but not in hypomorphic limb regeneration in *Xenopus laevis*

**DOI:** 10.1101/2022.12.27.522067

**Authors:** Reimi Tada, Takuya Higashidate, Takanori Amano, Shoma Ishikawa, Chifuyu Yokoyama, Saki Nara, Koshiro Ishida, Akane Kawaguchi, Haruki Ochi, Hajime Ogino, Nayuta Yakushiji-Kaminatsui, Joe Sakamoto, Yasuhiro Kamei, Koji Tamura, Hitoshi Yokoyama

## Abstract

*Xenopus* young tadpoles regenerate a limb with the anteroposterior (AP) pattern, but metamorphosed froglets regenerate a hypomorphic limb after amputation. The key gene for AP patterning, *shh*, is expressed in a regenerating limb of the tadpole but not in that of the froglet. Genomic DNA in the *shh* limb-specific enhancer, MFCS1 (ZRS), is hypermethylated in froglets but hypomethylated in tadpoles: *shh* expression may be controlled by epigenetic regulation of MFCS1. Is MFCS1 specifically activated for regenerating the AP-patterned limb? We generated transgenic *Xenopus laevis* lines that visualize the MFCS1 enhancer activity with a GFP reporter. The transgenic tadpoles showed GFP expression in *hoxd13-* and *shh-*expressing domains of developing and regenerating limbs, whereas the froglets showed no GFP expression in the regenerating limbs despite having *hoxd13* expression. Genome sequence analysis and co-transfection assays using cultured cells revealed that Hoxd13 can activate *Xenopus* MFCS1. These results suggest that MFCS1 activation correlates with regeneration of AP-patterned limbs and that re-activation of epigenetically inactivated MFCS1 would be crucial to confer the ability to non-regenerative animals for regenerating a properly patterned limb.

## INTRODUCTION

Among tetrapods, only urodele amphibians (newts and salamanders) can regenerate an amputated limb throughout their life cycles (Han et al., 2005; Joven et al., 2019; Otsuki and Tanaka, 2022). In *Xenopus laevis*, a tadpole can regenerate its limbs prior to metamorphosis, but it loses regenerative capacity gradually as metamorphosis proceeds (Dent, 1962; Muneoka et al., 1986). The froglet, a young post-metamorphosis adult, regenerates only a spike-like structure after limb amputation (Dent, 1962; reviewed in Suzuki et al., 2006): the spike lacks the anteroposterior (AP) pattern. In the limb formation process, *sonic hedgehog* (*shh*) is expressed posteriorly in the limb bud and serves as the key gene for the AP axis formation (Riddle et al., 1993; Harfe et al., 2004). *shh* is also expressed in the posterior part of a regenerating limb (blastema) in urodeles (Imokawa and Yoshizato, 1997; Torok et al., 1999). In *Xenopu*s, *shh* is expressed posteriorly in the limb blastema of a tadpole (Endo et al., 1997) but is not expressed in the blastema of a froglet limb (Endo et al., 2000; Yakushiji et al., 2007). Treatment with a Hedgehog inhibitor results in hypomorphic limb regeneration in both a *Xenopus* tadpole and an axolotl (Roy and Gardiner, 2002; Satoh et al., 2006). These results suggest that *shh* re-expression correlates with the limb regenerative capacity: whether a limb regenerates with the AP pattern or not.

Genetic studies in mice have revealed that a long-range enhancer called MFCS1 (mammal-fish-conserved-sequence 1), also called ZRS or Rr29, regulates limb-specific *Shh* expression (Lettice et al., 2003; Sagai et al., 2004). In transgenic (Tg) mouse, MFCS1 with a minimal promoter drives *lacZ* expression in the posterior part of a limb bud (Lettice et al., 2003). Depletion of a large part of MFCS1 results in limb-specific loss of *shh* expression and in loss of the AP polarity of a limb (Sagai et al., 2005). The activity of MFCS1 is positively regulated by Hoxd13 and ETS transcription factors, downstream factors of FGF signaling in a mouse limb bud that directly bind to MFCS1 (Capellini et al., 2006; Lettice et al., 2012). In amphibian limbs, the methylation level of genomic DNA in MFCS1 is high in the limbs of a *Xenopus* froglet, while the level is low to medium in the limbs of a *Xenopus* tadpole, an axolotl and a Japanese newt (Yakushiji et al., 2007). These findings strongly suggest that epigenetic inactivation of MFCS1 leads to the shutdown of *shh* expression that results in hypomorhic limb regeneration without the AP pattern in a froglet. The importance of MFCS1 for limb regeneration is further supported by the partial deletion of MFCS1 in an Iberian ribbed newt: it resulted in reduction in *shh* expression (Suzuki et al., 2018). However, the spatio-temporal activation pattern of MFCS1 has not been visualized in regenerating limbs of amphibians.

Here, we succeeded in visualizing the spatio-temporal pattern of MFCS1 activation in regenerating limbs by establishing Tg *Xenopus laevis* lines carrying a GFP linked to MFCS1 (MFCS1-GFP). Expression patterns of MFCS1-GFP were overlapped with those of *shh* both in limb development and limb regeneration. MFCS1-GFP expression was specifically localized in the regenerating limb of a young tadpole but was not found in the regenerating froglet limb. We revealed that the DNA methylation level of exogenous MFCS1 of the transgene as well as that of endogenous MFCS1 in the limb were higher in the froglet than in the tadpole. Our results suggest that MFCS1 is activated in limb regeneration with the AP pattern but is inactivated in hypomorphic limb regeneration after metamorphosis when MFCS1 has a higher DNA methylation status than that in tadpole stages.

## RESULTS AND DISCUSSION

### MFCS1-GFP expression localizes in the posterior mesenchyme of a developing limb bud

To visualize the MFCS1 activity, we generated a transgene construct containing GFP downstream of a *Xenopus shh* promoter (Yakushiji et al., 2007) and MFCS1 sequences (Fig. 1A). We generated transgenic *X. laevis* containing this transgene named “*Xla.Tg(Xtr.shh:GFP)^Tamura^*” (according to the Xenbase nomenclature). For highlighting the involvement of MFCS1 enhancer, we hereafter refer to this Tg as MFCS1-GFP. We observed the GFP pattern in G0 founders. Since *shh* is most prominently expressed posteriorly in a limb bud at stage 51-52 (Endo et al., 1997; Fig. 1B), we firstly observed at stage 51-52 (Nieuwkoop and Faber, 1994). GFP was found in the posterior part of the limb bud in at least eleven independent G0 individuals and the fluorescence gradually diminished after stage 52. Thus, we considered that MFCS1-GFP expression reflects the activity of the MFCS1 enhancer. We established two independent lines by mating selected GFP-expressing founder animals with wild-type frogs (Table S1). About 50% of the N1 individuals of line 2 as well as the N1 and N2 individuals of line 1 were GFP-positive in limb buds, presumably being inherited in a Mendelian manner. Limb buds of both line 1 and line 2 at stage 51-52 also showed GFP expression in the posterior part (Fig. 1C, C’ and Fig. S1A, A’). In the following experiments, we used GFP-positive individuals of line 1 (N1 or N2) and line 2 (N1).

**Fig. 1.**
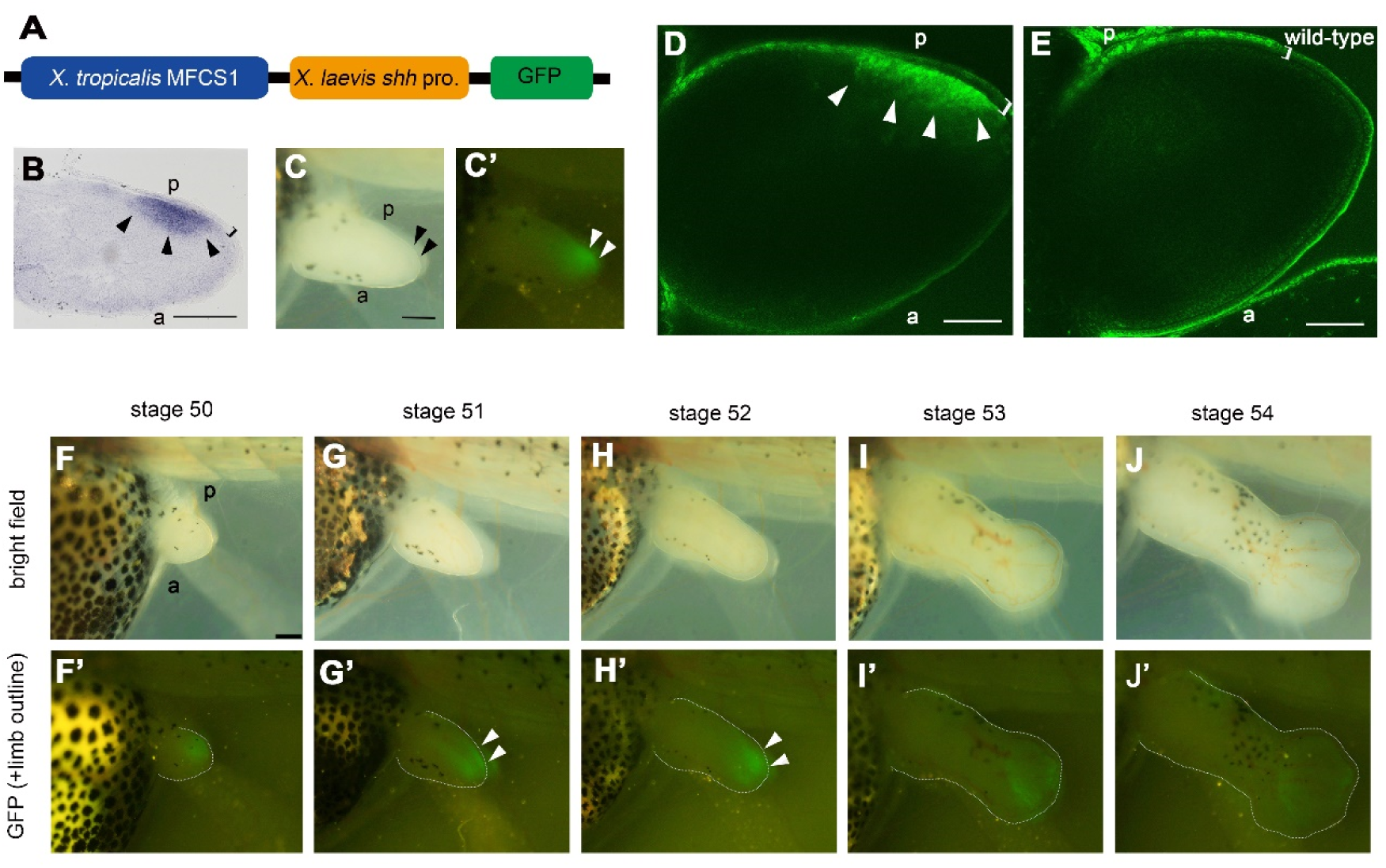
MFCS1-GFP expression coincides with *shh* expression in a developing limb bud. (A) The MFCS1-GFP transgene. (B) *shh* expression at stage 52. (C, C’) A limb bud of an MFCS1-GFP tadpole at stage 51. Bright-field (C) and fluorescent (C’) images. (D, E) Optical-section images by confocal microscopy of the limb bud of an MFCS1-GFP (D) and that of a wild-type tadpole at stage 51. Bright-field (F-J) and fluorescent (F’-J’) images of the limb bud of the same MFCS1-GFP tadpole. Limb buds are outlined by dotted lines in the fluorescent images. All of the MFCS1-GFP tadpoles shown are line 1. a, anterior: p, posterior. Arrowheads, *shh* (B) or GFP expression (D, E, G’, H’) at the posterior side: Bracket, the epidermal cell layer. Bars = 100 μm for (B, D, E) and 200 μm for (C, C’, F-J, F’-J’).

Confocal imaging revealed GFP localization in the posterior mesenchyme of the limb bud at stage 51-52 (Fig. 1D, E and Fig. S1B, C). In limb development, GFP was firstly observed at stage 50 in a limb bud (Fig. 1F, F’ and Fig. S1D, D’). This observation is consistent with the initial weak *shh* expression detected at stage 50 by *in situ* hybridization (Endo et al., 1997). GFP expression became strongest at stages 51-52 and the fluorescence was more posteriorly polarized by stage 52 (Fig. 1G, G’, H, H’ and Fig. S1E, E’, F, F’). At stage 53, the fluorescence was weaker than that at stage 52 (Fig. 1I, I’ and Fig. S1G, G’). Finally GFP expression was almost undetectable at stage 54 (Fig. 1J, J’ and Fig. S1H, H’). These observations were, in principle, consistent with the spatio-temporal *shh* expression patterns (Endo et al., 1997), while the GFP expression appeared to be broader than that of *shh* expression. We concluded that the MFCS1-GFP expression pattern in the Tg lines mostly visualizes the activity of the endogenous MFCS1 enhancer in a developing limb bud.

### MFCS1-GFP expression predicts the AP duplication of a limb bud

To further examine whether MFCS1-GFP expression correlates with limb morphogenesis along the AP axis, we conducted a 180° rotation experiment on limb buds of Tg (line 1) tadpoles (Fig. 2A). Cameron and Fallon (1977) found limb duplication along the AP axis after rotation of the distal limb bud tips 180° in *X. laevis*. Endo et al. (1997) repeated this experiment and found that ectopic *shh* expression was induced in the posterior region of the graft and stump. To test the correlation between MFCS1 enhancer activity and limb duplication, we repeated this experiment (Fig. 2). Consistent with the preceding results in control limb buds without rotation, showing only faint ectopic *shh* expression (Endo et al., 1997), the control limb buds (ten of 12 samples) did not show GFP expression and did not result in limb duplication (Fig. 2B, C). In contrast, the 180° rotated limb buds (eight of 11 samples) showed ectopic induction of GFP expression in the posterior part of the graft and finally formed duplicated limbs along the AP axis (Fig. 2D-G). These results (summarized in Table S2) suggest that ectopic induction of MFCS1 activity is closely related to ectopic *shh* expression and subsequent limb duplication along the AP axis, further suggesting that MFCS1-GFP expression reflects the endogenous *shh* enhancer activity.

**Fig. 2.**
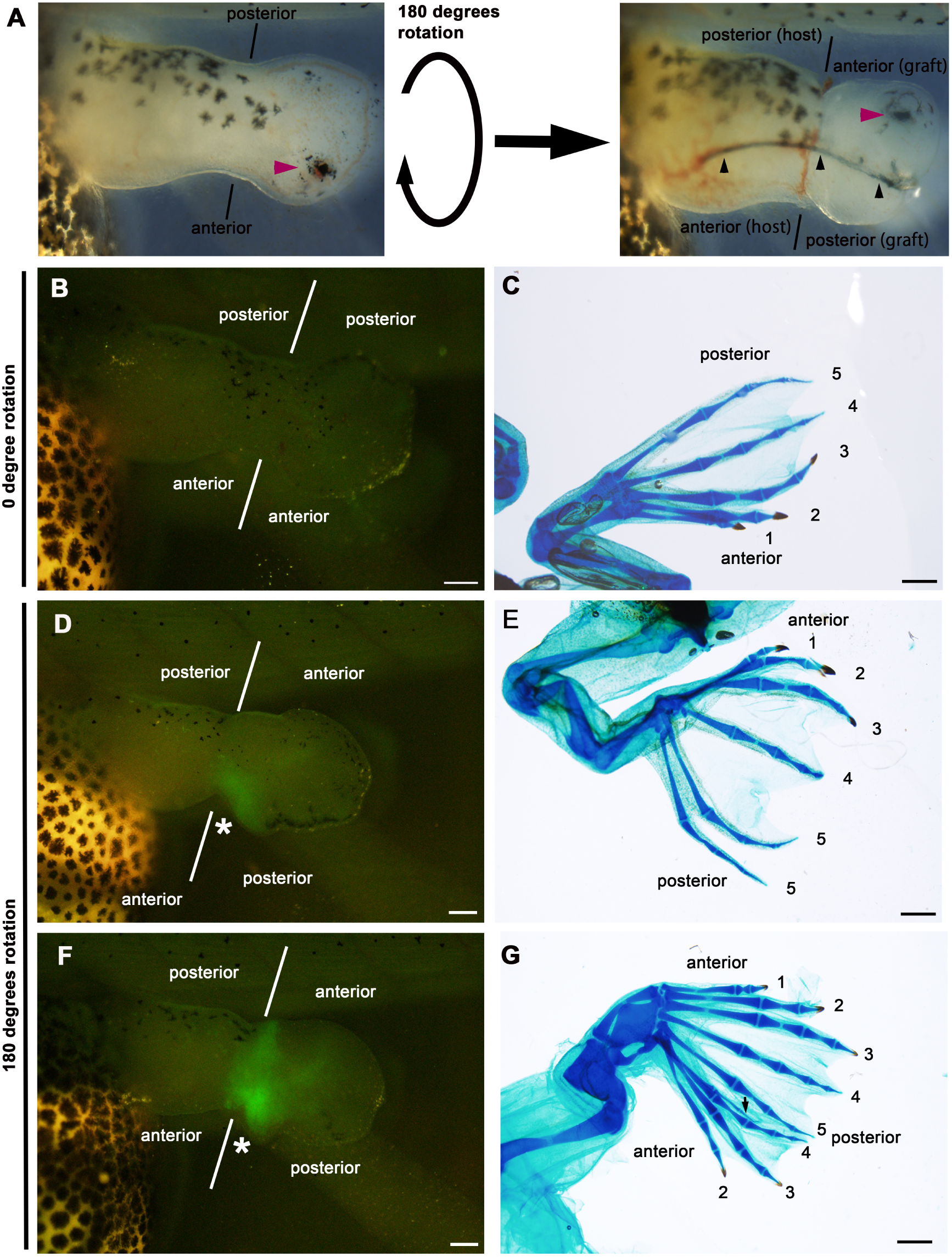
MFCS1-GFP expression precedes limb duplication along the AP axis in 180° rotated limb buds. (A) 180° rotation of a limb bud at stage 53. A magenta arrowhead, a Nile blue particle: Black arrowheads, a tungsten pin. (B, C) A 0° rotated limb bud at 6 days after grafting (B) and the resultant normal limb formation (C). (D-G) 180° rotated limb buds. Rotated limb buds at 6 days after grafting (D,F) resulted in supernumerary limb formation (E,G). A pair of lines, the host-graft boundary: An asterisk, ectopic GFP: An Arabic number, the digit identity: A black arrow, an unidentifiable digit. Bars = 200 μm for (B, D, F) and 1 mm for (C, E, F).

### MFCS1 is activated in a blastema that regenerates a limb with the AP pattern

To examine whether MFCS1 is activated in limb regeneration with the AP pattern, we amputated a stage 54 limb bud at the ankle level: this amputation results in almost complete limb regeneration (Muneoka et al., 1986). By this stage, *shh* and MFCS1-GFP expression in a limb bud have become almost undetectable (Fig. 1 and Fig. S1) (Endo et al., 1997). At 4 days post amputation (dpa), GFP was observed broadly in the regenerating blastema (Fig. 3R and Fig. S2B-B’’), indicating that MFCS1 is re-activated in the blastema that regenerates a well-patterned limb.

**Fig. 3.**
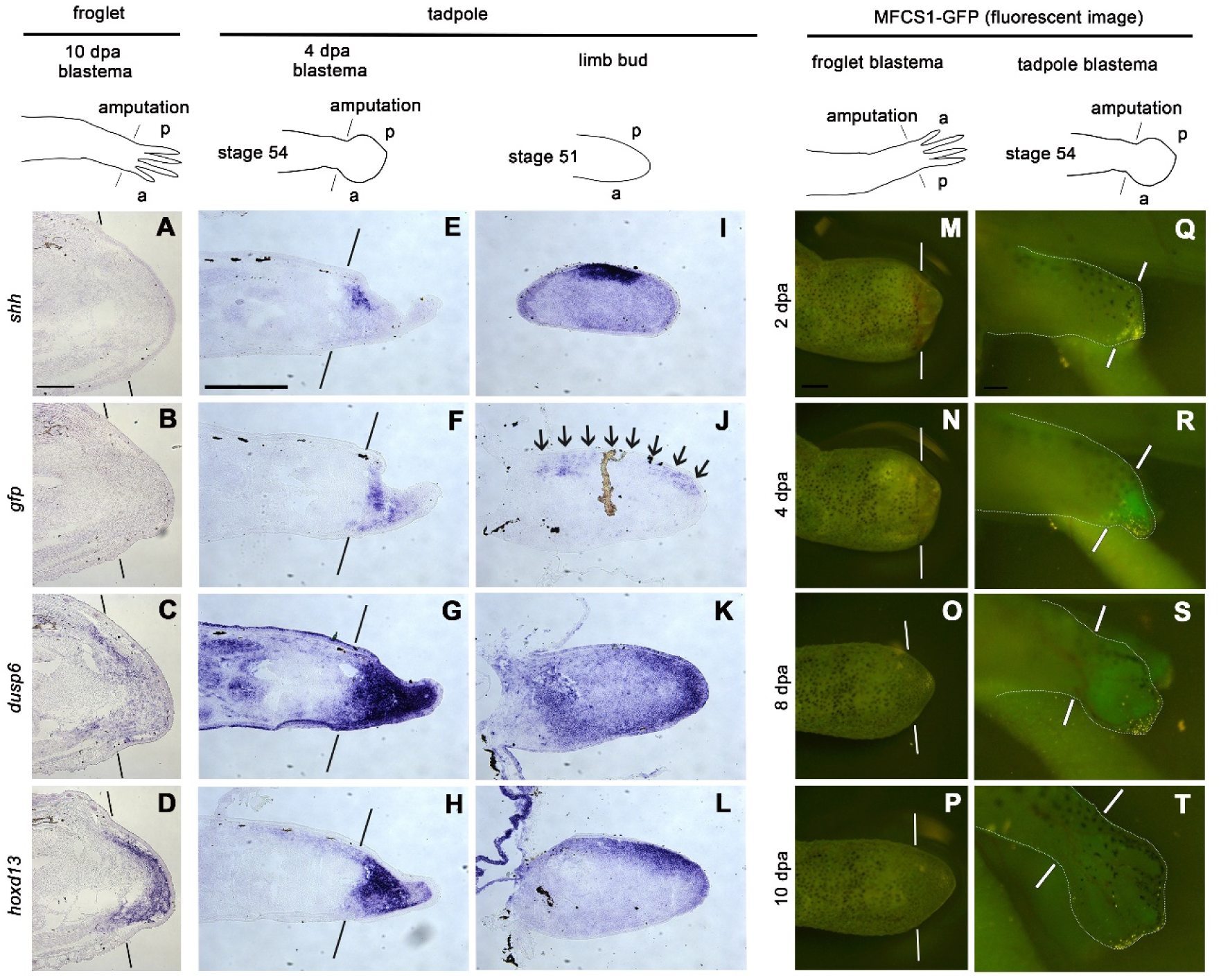
Comparison of gene expression patterns and MFCS1-GFP expression pattern in a regenerating limb in a young tadpole and a froglet. (A-D) Adjacent sections of a blastema of a froglet. n=5 of 5 blastemas (line 1). (E-H) Adjacent sections of a blastema of a tadpole. n=9 of 9 (line 1) and 6 of 6 blastemas (line 2), respectively. (I-L) Adjacent sections of a limb bud. n=10 of 10 (line 1) and 6 of 6 limb buds (line 2), respectively. (M-P) A blastema of a froglet of line 1. See also Fig. S3. (Q-T) A blastema of a tadpole of line 1. The blastema and the stump are outlined by a dotted line. See also Fig. S2. a, anterior: p, posterior. A pair of lines, the amputation level: Arrows, *gfp* expression at the posterior side. All pictures shown in Fig. 3 are samples of line 1. Bar = 200 μm for (A-L, Q-T) and 500 μm for (M-P).

MFCS1-GFP expression was distributed throughout the blastema and not localized posteriorly. It became stronger at 6 dpa (Fig. S2C-C’’), then the expression gradually diminished (Fig. 3S, T and Fig. S2). *shh* expression was restricted posteriorly in a blastema at 4 dpa (Fig. 3E). Based on these results, we judged that MFCS1-GFP expression is distributed broadly in regenerating blastemas and that the temporal expression pattern of MFCS1-GFP coincidences with that of *shh* expression, while the spatial MFCS1-GFP expression pattern is wider than the *shh-*expressing domain. MFCS1-GFP signal in the regenerating blastema of a young tadpole suggests that MFCS1 is re-activated in a regenerating limb with the AP pattern formation when *shh* is expressed: *shh* expression in the regenerating limb as well as in the developing limb seems to be regulated by the common enhancer, MFCS1.

### MFCS1-GFP expression pattern correlates with the pattern of *hoxd13* in a tadpole

Hoxd13 and FGF downstream factors, ETS transcription factors, positively regulate the MFCS1 enhancer in mice (Capellini et al., 2006; Lettice et al., 2012). We examined *dusp6* (*mkp3*) and *hoxd13* expression as indicators of positive upstream regulators of MFCS1. *dusp6* expression is induced by FGF signaling (Eblaghie et al., 2003; Kawakami et al., 2003) and especially reflects the activity of ETS transcription factors (Ekerot et al., 2008). We chose a stage 51 limb bud and a 4 dpa blastema of a tadpole in which the limb bud was amputated at the ankle level at stage 54 as representatives of developing and regenerating limbs, respectively: both initiate prominent *shh* expression in the posterior side (Fig. 3E, I). In a developing limb bud, *dusp6* is expressed broadly and strong expression is seen especially in the apical mesenchyme (Fig. 3K). *hoxd13* is expressed in the posterior mesenchyme (Fig. 3L) and its expression domain is wider than the *shh*-expressing domain (Fig. 3I). The posterior mesenchymal region of a limb bud is positive for both *dusp6* and *hoxd13* and the double positive region coincides with the fluorescence of MFCS1-GFP (Fig. 1C’, D, G’) and *gfp* transcript (Fig. 3J) patterns. The *gfp* transcript was not detected in line 2 limb bud samples (n=0/6) but was detected in line 1 limb bud samples (n=10/10) by *in situ* hybridization, suggesting that line 1 has stronger transgene expression than that of line 2. The coincidence between upstream regulators and MFCS1-GFP expression was clearer in a regenerating limb: *dusp6* (Fig. 3G) and *hoxd13* (Fig. 3H) were strongly expressed in whole region of a blastema. MFCS1-GFP expression (Fig. 3R) and *gfp* transcript (Fig. 3F) are robustly distributed in the entire blastema, while *shh* expression is restricted posteriorly (Fig. 3E). Therefore, the MFCS1-GFP expression domain is larger than the *shh* expression domain, especially in a regenerating limb. Transcriptional repression by a silencer or defined regulation of chromatin conformational change resulting in the proximity of MFCS1 and the *shh* promoter (Amano et al., 2009) may further restrict endogenous *shh* expression to the posterior edge of the blastema. Even though *shh* expression is spatially narrower than the GFP expression pattern, the expression pattern of MFCS1-GFP covers the *shh*-expressing domain and coincides with the expression domains of upstream regulators represented by *hoxd13* and *dusp6*.

### MFCS1 is inactive in a limb blastema of a metamorphosed froglet despite *hoxd13* expression

A limb of a metamorphosed froglet does not express *shh* after amputation, and amputation results in regeneration of a spike, while an amputated limb of a froglet forms a regeneration blastema (Endo et al., 2000; Suzuki et al., 2006; Yakushiji et al., 2007). Hypermethylated genomic DNA at the MFCS1 strongly suggested that this enhancer is inactivated in a froglet limb: it implies transcriptional inactivation of *shh* in the limb (Yakushiji et al., 2007). To examine whether inactivation of the enhancer occurs after metamorphosis, we examined MFCS1-GFP expression in limb regeneration of a froglet. From the early to the late stage, no specific GFP was observed in the blastema (Fig. 3M-P and Fig. S3). Therefore, we conclude that MFCS1 is inactivated completely or is much more inactive in the blastema of a froglet limb than in that of a young tadpole.

We then examined expression of upstream regulators of MFCS1. By 10 dpa, an amputated limb of a froglet forms a cone-shaped blastema and it already starts expressing *fgf-8*, *fgf-10*, *msx1*, and *hand2* (Suzuki et al., 2005; Yakushiji et al., 2007). In addition, Yakushiji et al. (2007) reported hypermethylation of genomic DNA at the MFCS1 in a limb blastema at 10 dpa in a froglet. Therefore, we chose the blastema at 10 dpa and examined gene expression. At 10 dpa, *hoxd13* was broadly expressed (Fig. 3D). *dusp6* was also expressed (Fig. 3C), while no *shh* (and *gfp*) expression was detected in the blastema as Yakushiji et al. (2007) reported (Fig. 3A, B). The expression of *hoxd13* and *dusp6* in a froglet blastema suggests that upstream regulators of MFCS1 (5’ Hoxd and ETS transcription factors) as well as *hand2* (Yakushiji et al., 2007) are not absent in the froglet blastema. Thus, MFCS1 in a froglet limb appears to be unable to respond to positive regulators represented by Hoxd13.

### Higher DNA methylation level of MFCS1 may inactivate the enhancer’s responsiveness to Hoxd13 in a froglet limb

A local alignment of the highly conserved sequences revealed multiple putative Hox-binding motifs in MFCS1 (Fig. 4A, see also Fig. S4). To confirm the potential responsiveness of MFCS1, MFCS1-*shh* promoter-*luc2* or MFCS1-minP-*luc2* reporter constructs (Fig. 4A) were co-transfected with the gene expression constructs into NIH3T3 cells. The reporter activities with MFCS1-*shh* promoter-*luc2* were significantly increased with *Fgf2* (1.4-fold vs. mock), *Fgf4* (1.5-fold vs. mock), *Fgf8* (1.3-fold vs. mock), or *Hoxd13* (2.3-fold vs. mock) (Fig. 4B). When the MFCS1-*luc2* was artificially methylated by CpG methyltransferase in advance, the responsiveness to Hoxd13 was strongly suppressed by ∼260 fold (Fig. S5). Taken together with the highly methylated MFCS1 in a *Xenopus* froglet limb (Yakushiji et al., 2007), this result raises the possibility that DNA methylation of MFCS1 inactivates its responsiveness to Hoxd13.

**Fig. 4.**
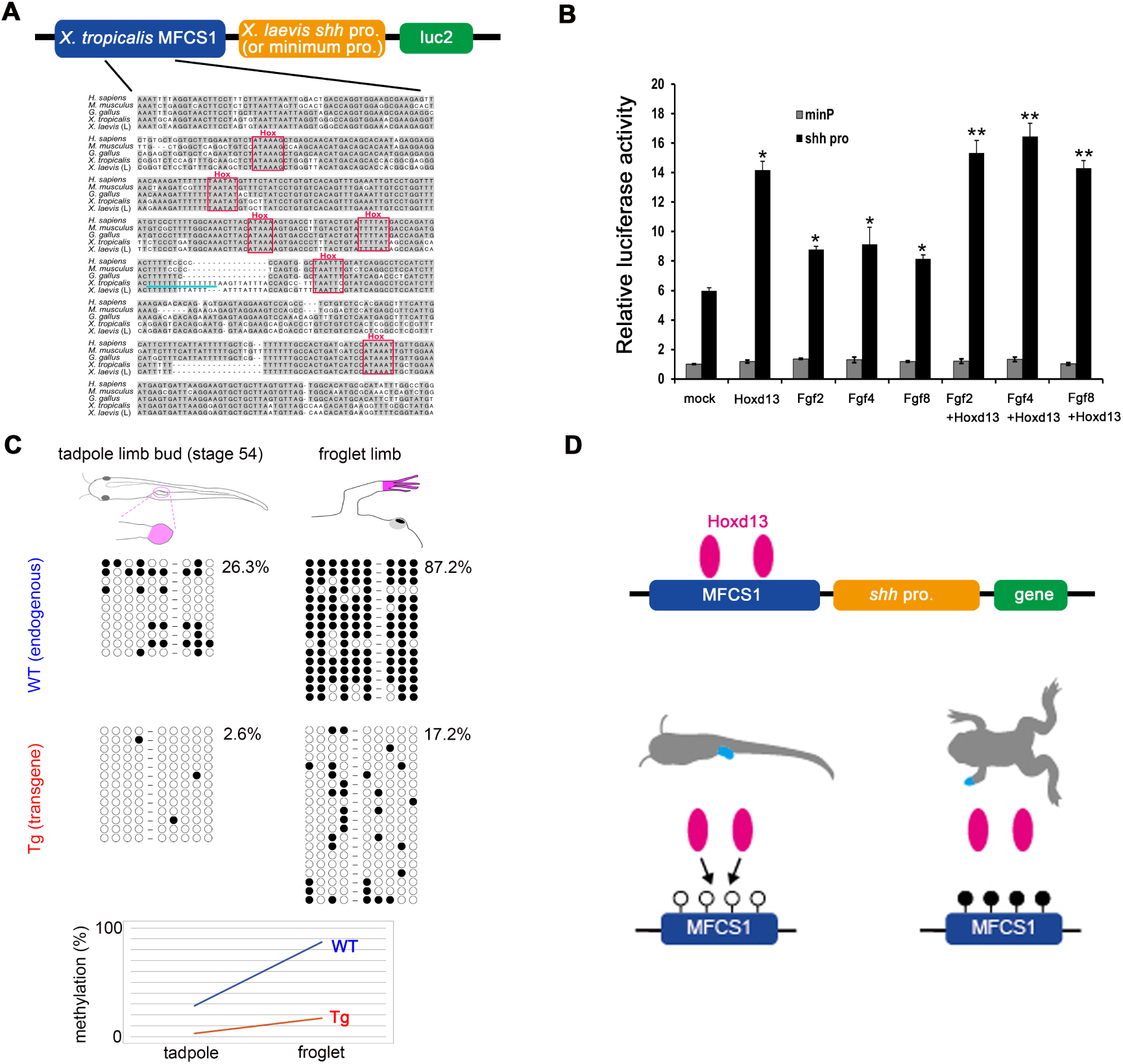
Transactivation and DNA methylation of *Xenopus* MFCS1 enhancer. (A) Upper: Map of MFCS1-*luc2* for assays in (B). Lower: Alignment of MFCS1 highly conserved sequences. Nucleotides identical in more than two sequences are shaded. Conserved Hox-binding motifs are boxed. A light bule line underlies the 14-thymidine repeat. (B) Normalized luciferase activity. Error bars, standard deviation from the mean (n=3). *Significant increase from mock (p<0.01). **Significant increase in Fgf2+Hoxd13, Fgf4+Hoxd13 and Fgf8+Hoxd13 from sole Fgf2, Fgf4 and Fgf8, respectively (p<0.01). (C) Methylation pattern of endogenous and transgene MFCS1 (line 1). Samples were obtained from the autopod (pink). Circles correspond to nine CpG sites (Fig. S6). Lower: Increases in average methylation rate of the MFCS1 after metamorphosis. (D) Model for epigenetic silencing of gene expression. Hoxd13 activates hypomethylated MFCS1 and results in transcription of a gene (left). However, Hoxd13 is unable to activate highly methylated MFCS1 and results in inactive transcription (right).

To further examine whether a high methylation level of MFCS1 exerts a suppressive effect on gene expression, we investigated the methylation status of MFCS1 as the transgene in MFCS1-GFP Tg tadpoles and froglets. Within an MFCS1 highly conserved sequence in amphibians, there are nine CpG sites in *X. tropicalis* and *X. laevis* (Fig. S6). Firstly, we examined DNA methylation status of MFCS1 in wild-type tadpoles and froglets by bisulfite sequencing as reported by Yakushiji et al. (2007) and obtained consistent results: Young tadpoles have a lower methylation (26.3%) and froglets have a higher methylation level (87.2%) in the limbs (Fig. 4C). We then examined the methylation level of MFCS1 as the transgene in MFCS1-GFP Tg (line 1). Again, froglet limbs have a higher methylation level (17.2%) than that in limb buds of young tadpoles (2.6%) (Fig. 4C). After limb amputation in a froglet and after completion of spike regeneration, a higher methylation level than that in tadpoles was still found in the Tg regenerates (line 1, 14.4%) (Fig. S7). MFCS1 as the transgene was susceptible to methylation to some extent also in line 2 (Fig. S7). Although the basal methylation level differs between endogenous MFCS1 and transgene MFCS1, the increase in the methylation level of the transgene MFCS1 after metamorphosis also correlated with the ontogenetic inactivation of gene expression under control of the enhancer.

## Conclusions

In this study, we visualized MFCS1 activity in limb regeneration of an amphibian. MFCS1 is re-activated in the regenerating blastema for a patterned limb in a young tadpole but not in the blastema for a hypomorphic limb in a froglet. MFCS1 seems to be activated by Hoxd13 in a tadpole but unable to respond to Hoxd13 in a froglet. An ontogenetic increase in the MFCS1 methylation may block the function of Hoxd13 (and other positive regulators) as a transcriptional activator(s) to the enhancer so that transcription of a gene under the enhancer is inactivated (Fig. 4D). For improvement of regenerative capacity in a froglet limb by re-activation of *shh* expression, removal of epigenetic silencing of MFCS1 would be important. The combinational application of an inhibitor of DNA methyltransferase and a histone deacetylase inhibitor stimulated *shh* expression in cultured limb blastema cells of a froglet (Yakushiji et al., 2009), suggesting that application of such reagents could be hopeful if the condition is optimized *in vivo*. As an alternative approach, CRISPR-based epigenome editing that removes DNA methylation (e.g., Nuñez et al., 2021) may be effective. In these approaches, the MFCS1-GFP Tg line would be useful for evaluation: The enhancer activation is easily detected by GFP and it would thus be possible to screen chemicals that remove epigenetic silencing of the enhancer in a large scale. If regeneration of a well-patterned limb is induced in a froglet by re-activation of MFCS1, it would be a precious model case for recovery of organ-level regeneration in non-regenerative higher vertebrates by manipulating epigenetics.

## MATERIALS AND METHODS

### Transgene construction and *in situ* hybridization

A 3,164-bp MFCS1 enhancer sequence was amplified from *Xenopus tropicalis* genomic DNA prepared from a Nigerian line with primers (XtMFCS1-Fw and Xt-MFCS1-Rv; Table S3). The PCR product was cloned into the pCRII-TOPO vector (Invitrogen) and sequenced. The *Xenopus laevis shh* promoter (Yakushiji et al., 2007) was cloned into a reporter construct that contains an EGFP expression cassette. Then the MFCS1-*shh* promoter-*egfp* was generated by inserting the 3,164-bp MFCS1 enhancer sequence upstream of the *shh* promoter (Fig. 1A). For synthesis of probes for *in situ* hybridization, a *hoxd13* cDNA clone, BC080434, inserted in pCMV-SPORT6 was obtained from Dharmacon. The *hoxd13*/ pCMV-SPORT6 plasmid was linearized with SalI and transcribed with T7 RNA polymerase. DIG-labeled RNA probes of *hoxd13*, *shh* (Endo et al., 1997), *gfp* (Hayashi et al., 2015) and *dusp6* (Gomez et al., 2005) were prepared according to the protocol of the manufacturer (Roche). To prepare serial cryosections, the specimens were fixed in MEMFA (0.1 M MOPS at pH 7.4, 2 mM EGTA, 1 mM MgSO_4_, 3.7% formaldehyde) overnight at room temperature, embedded in OCT compound (Sakura), and serially sectioned at 8-10 µ m by a cryostat. *In situ* hybridization on frozen sections was performed according to Yoshida et al. (1996) with slight modifications. Sections were observed under an upright microscope (Olympus BX53) with an objective lens (Olympus UPlan PLN 10X) and were photographed by a digital camera (Leica DFC7000T).

### Animal husbandry and transgenesis in *Xenopus laevis*

Wild-type *Xenopus laevis* adult frogs were purchased from domestic animal venders, Watanabe Zoushoku (http://www5d.biglobe.ne.jp/~zoushoku/top.htm) and Hamamatsu Seibutsu Kyozai (http://www.h-seibutsu.co.jp/index.html). Founder (G0) MFCS1-GFP Tg *Xenopus laevis* embryos were generated by a REMI technique (Kroll and Amaya, 1996) using oocyte extract instead of egg extract (Hirsch et al., 2002; reviewed in Ogino and Ochi, 2009). MFCS1-GFP N1 Tg *Xenopus laevis* lines were obtained by crossing sexually mature MFCS1-GFP G0 frogs with wild-type frogs. Tadpoles and froglets were reared at 22–23°C in dechlorinated tap water. The rearing containers were cleaned daily, and the tadpoles were fed powdered barley grass (Odani Kokufun, Japan or Yamamoto Kanpoh, Japan). At stage 58 (Nieuwkoop and Faber, 1994), the feeding was ceased until metamorphosis was completed. After metamorphosis, the froglets were fed dried tubifex every other day. All surgery was performed under ethyl-3-aminobenzoate anesthesia, and all efforts were made to minimize suffering. When tadpoles and froglets needed to be euthanized, they were immersed in 0.05% ethyl-3-aminobenzoate (Tokyo Chemical Industry, 886-86-2). When adult male frogs were sacrificed to excise their testes for transgenesis, 2% ethyl-3-aminobenzoate dissolved in pure water was injected into the intracardiac cavity (400 µ g of ethyl-3-aminobenzoate/1 g of animal). After the male frog was confirmed to be completely unconscious, its testes were excised with a blade, and then the unconscious frog was put in a freezer for euthanization. All animal care and experimentation procedures were conducted in accordance with the animal care and use committee guidelines of Tohoku University (2015LsA-023) and Hirosaki University (A15003, A15003-1).

### Observation of developing and regenerating limbs, limb amputation and 180° rotation of a limb bud

For observation of limb development and limb regeneration in tadpoles, we used hindlimb buds since it is much easier to observe and to operate on hindlimb buds than forelimb buds due to their size and accessibility. To observe the regenerating limb bud (blastema) of a tadpole, tadpoles at stage 54 were anesthetized with 0.025% ethyl-3-aminobenzoate and hindlimb buds were amputated using ophthalmologic scissors at the presumptive ankle level (according to the outside view and a fate map by Tschumi, 1957). For preparation of a 180° rotated limb bud, Nile Blue A particles were embedded into the anterior side of the distal tip of a stage 53 hindlimb bud with a tungsten needle as the reference point indicating the original AP axis of the graft according to Cameron and Fallon (1977). Subsequently, the limb bud was amputated at the presumptive ankle level (Tschumi, 1957) and then the distal limb bud tip was rotated 180° (not rotated for control limbs) on the proximo-distal axis, returned to the stump, and fixed with tungsten pins according to Cameron and Fallon (1977) and Endo et al. (1997). After metamorphosis, the resulting limbs were fixed in 10% formalin in Holtfreter’s solution and stained with 0.1% Alcian blue as previously reported (Yokoyama et al., 1998). Then digit identity was determined by the number of phalanges in association with the presence/absence of a claw (Cameron and Fallon, 1977). To observe the blastema of a froglet, froglets were anesthetized with 0.05% ethyl-3-aminobenzoate and then forelimbs were amputated through the distal zeugopodium with ophthalmologic scissors and the amputation surface was trimmed to be flat. For observation of limb regeneration in froglets, we used forelimbs since hindlimb amputation often results in drowning of the froglets.

### Imaging of whole-mount limb samples

Tadpoles and froglets anesthetized with 0.025% or 0.05% ethyl-3-aminobenzoate were observed under a fluorescence stereomicroscope (Nikon SMZ18) with a 0.5x objective lens using a GFP longpass filter (if in a fluorescent view) and photographed by a digital camera (Olympus DP74 or DP73). For confocal microscopy (Fig. 1D, E and Fig. S1B, C), anesthetized tadpoles were each placed on a 35-mm glass-bottom dish (Iwaki, Japan) so that the dorsal surface of the hindlimb bud was tightly attached to the glass-bottom. The prepared samples were observed under an FV3000 confocal laser scanning microscope (Olympus) equipped with an Olympus UplanSApo 20× dry objective lens. Fluorescence between 500 nm and 600 nm was detected by a GaAsP photo multiplier tube. Optical-section images at approximately 60-80 μm in depth from the dorsal surface of the limb bud were photographed.

### Genome sequence analysis

The 449-b genomic sequences corresponding to a conserved core region of human MFCS1 (hg38, chr7: 156,791,242–156,791,690) and its orthologous sequences in the mouse (mm39, chr5: 29,520,033–29,520,474), chicken (galGal6, chr2: 8,553,351– 8,553,801), *X. tropicalis* (xenTro10, chr6: 17,346,249–17,346,700) and *X. laevis* (J-strain 10.1, Chr6L: 15,900,483–15,900,931) were downloaded from the UCSC Genome Browser and Xenbase (Karolchik et al., 2003; Fortriede et al., 2020). These sequences were aligned using MUSCLE (Edgar, 2004). Searches for potential transcription factor-binding sites were performed with transcription factor-binding motifs collected from the TRANSFAC and JASPAR databases (Wingender, 2008; Castro-Mondragon et al., 2022). The *X. tropicalis* MFCS1 element used for our functional analyses contains a 12-thymidine repeat in place of the 14-thymidine repeat underlined in the alignment (Fig. 4A, underlined). This difference may be because of a polymorphism between the strain used for our study and that used for the genome sequencing project (Igawa et al., 2015).

### Reporter construction, cell culture and luciferase assay

The highly conserved 998-bp core sequence of *Xenopus tropicalis* MFCS1 was amplified by PCR with specific primers (Table S3) and inserted into the Acc65I and XhoI sites of the pGL4.23 vector (Promega, E8411). Then the minimal promoter of the pGL4.23 vector was replaced with a 486-bp sequence of *Xenopus laevis shh* promoter (Fig. 4A). It should be noted that the 14-thymidine repeat in *X. tropicalis* MFCS1 sequence from the UCSC Genome Browser (Fig. 4A, underlined) was shortened to a 12-thymidine repeat in the transgene construct (Fig. S6) and the reporter construct for the luc assay. pGL4.74 containing the *Renilla* luciferase gene was used for normalization of reporter expression (Promega, E6921). Coding sequences of mouse *Hoxd13*, *Fgf2*, *Fgf4* and *Fgf8* were cloned into the pcDNA3.1 vector (ThermoFisher, V79020). NIH3T3 cell line (RCB2767) was obtained from the RIKEN BioResource Research Center. NIH3T3 cells were maintained in Dulbecco’s modified Eagle’s medium (ThermoFisher, 11960-044) containing 10% fetal bovine serum, 1x GlutaMax and 100 μg/ml penicillin–streptomycin. Cells at 70– 90% confluency were transfected with equal amounts of luciferase reporter and gene expression plasmids using Lipofectamine 3000 (ThermoFisher). Luciferase activity in the cell lysates was measured using a Dual-Luciferase Reporter Assay System (Promega) according to the manufacturer’s protocol. Firefly luciferase activity of MFCS1-*luc2* using a minimum promoter (minP) or *shh* promoter (shh pro) was normalized to *Renilla* luciferase control activity. The mean value of mock samples by the control construct with the minimum promoter was adjusted to 1.0. For statistical analysis, Student’s *t*-tests (two-tailed) were applied when equal variance tests were passed.

### Bisulfite genomic sequencing

Presumptive autopods of hindlimb buds (Tschumi, 1957) from thirty-five wild-type and sixteen MFCS1-GFP Tg (line 1) tadpoles at stage 54 were collected. Forelimb autopods from five wild-type and six MFCS1-GFP Tg (line 1) froglets were collected. Spike regenerates from two MFCS1-GFP Tg (line 1) and two same Tg (line 2) frogs were collected. Then genomic DNA was extracted from these tissue samples. The genomic DNA was treated with sodium bisulfite using an EZ DNA Methylation-Gold Kit following the protocol of the manufacturer (Zymo Research). DNA was amplified by 30 cycles of a three-step PCR (98℃ for 10 sec, 60℃ for 30 sec and 68℃ for 30 sec) with MFCS1_BS_Fw and MFCS1_BS_Rv primers (Table S3) using KOD -Multi & Epi-(Toyobo). Each product was cloned into the pANT vector (Nippon Gene) or pGEM-T easy (Promega) and sequenced with M13F-pUC or M13R-pUC primers (Table S3).

## ACKNOWLEDGMENTS

We thank Yoshiko Yoshizawa and Natsume Sagawa for excellent animal care. We thank Drs. Makoto Asashima, Shuji Takahashi and Yoshio Yaoita for providing the Nigerian line of *X. tropicalis*. We thank Dr. Ikuo Uchiyama for support in gene expression analysis. This work was partially supported by the NIBB Collaborative Research Program to HY (19-451, 20-436, 21-307, 21-401). This work was supported by the National Bio-Resource Project (NBRP) of the MEXT, Japan. We acknowledge Shared Facility Center for Science and Technology, Hirosaki University (SFCST) for confocal microscope observation by FV3000 (Olympus). We thank Dr. Chisato Ushida, Dr. Michiko Sasabe, Shunsuke Hosoi, Misako Saida and Hibiki Yokoyama for their help in using confocal microscopy. We thank Dr. Daisuke Kurita for help in TA cloning. We thank Drs. Gembu Abe and Kazuya Kobayashi for their help in preparation of tungsten needles. We thank Dr. Jose F. de Celis for providing *Xenopus laevis dusp6* cDNA.

## Author contributions

Conceptualization: K.T., H.Y.; Methodology: T.A., N.Y.-K., J.S., Y.K., H.Y.; Validation: T.A., N.Y.-K., H.Y.; Formal analysis: T.A., H.Ogino, H.Y.; Investigation: R.T., T.H., T.A., S.I., C.Y., S.N., K.I., A.K., H.Ochi, H.Ogino, N.Y.-K., Y.K., H.Y.; Resource: T.A., A.K., H.Ochi, H.Ogino, N.Y.-K., Y.K., K.T., H.Y.; Writing – original draft preparation: T.A., H.Ogino, J.S., Y.K., H.Y.; Writing – review and editing: A.K., H.Ochi, H.Ogino, N.Y.-K., K.T., H.Y.; Visualization: R.T., T.H., T.A., S.I., H.Ogino, N.Y.-K., H.Y.; Supervision: K.T., H.Y.; Project administration: H.Ogino, K.T., H.Y.; Funding acquisition: K.T., H.Y.

---

**Fig. S1.**
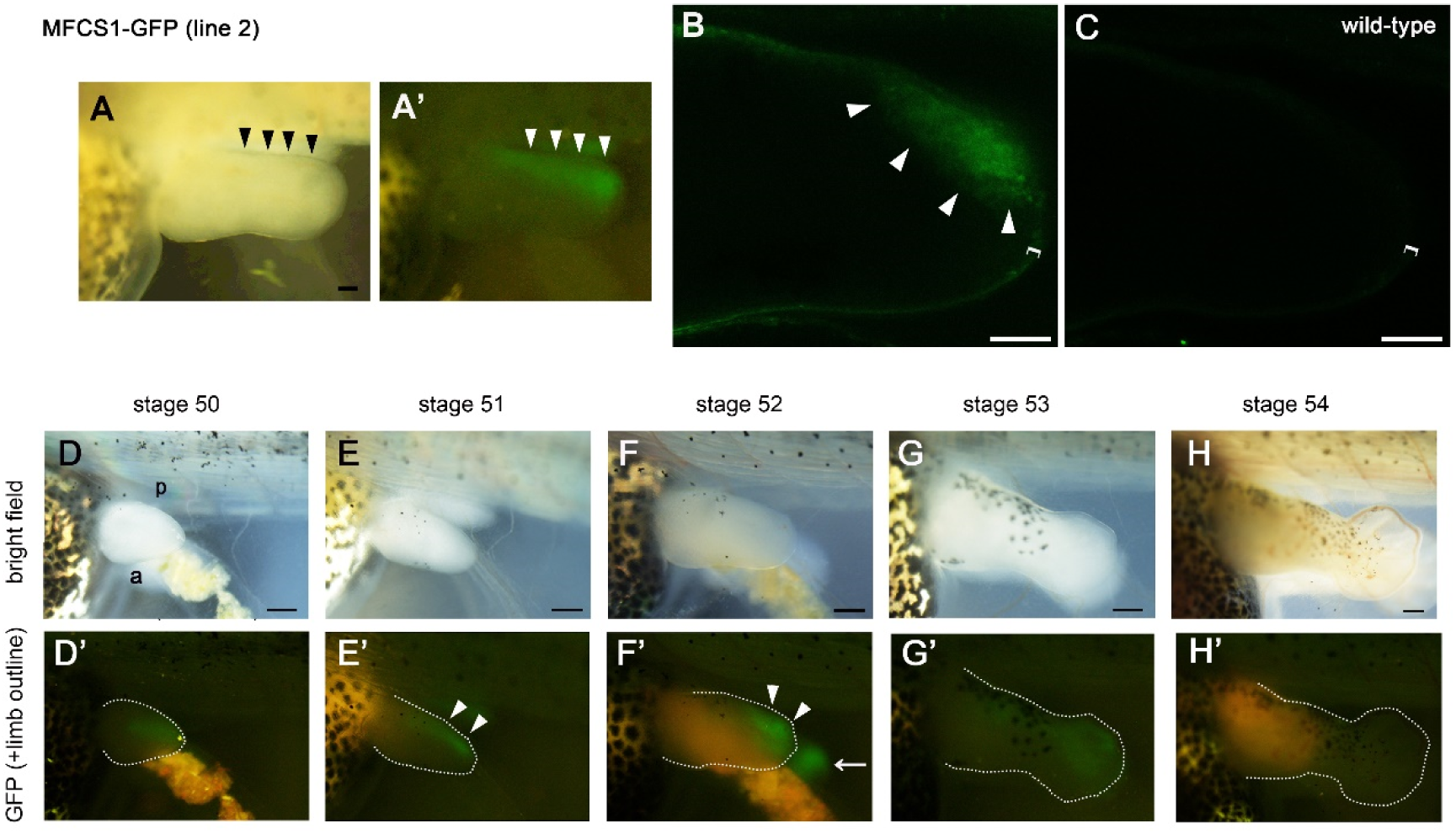
MFCS1-GFP expression pattern was recapitulated in line 2. (A) A developing limb bud of an MFCS1-GFP Tg tadpole of line 2 at stage 52. Bright-field (A) and fluorescent (A’) images of the same limb bud. (B, C) Optical-section images by confocal microscopy of the limb bud of an MFCS1-GFP Tg tadpole of line 2 (B) and a wild-type tadpole (C) at stage 52. GFP expression was detected in the posterior side mesenchyme specifically in the MFCS1-GFP Tg tadpole. Bright field (D-H) and fluorescent (D’-H’) images of the limb bud of the same Tg tadpole of line 2. Limb buds are outlined by dotted lines. GFP expression was detected in the posterior side of limb buds at stage 51-52 and then gradually diminished. a, anterior: p, posterior. Arrowheads, GFP expression (A, A’, B, E’, F’) at the posterior side of the limb buds: Bracket, the epidermal cell layer: A pair of lines, the amputation level. Bars = 100 μm for (A, B, C), 200 μm for (D-H, D’-H’).

**Fig. S2.**
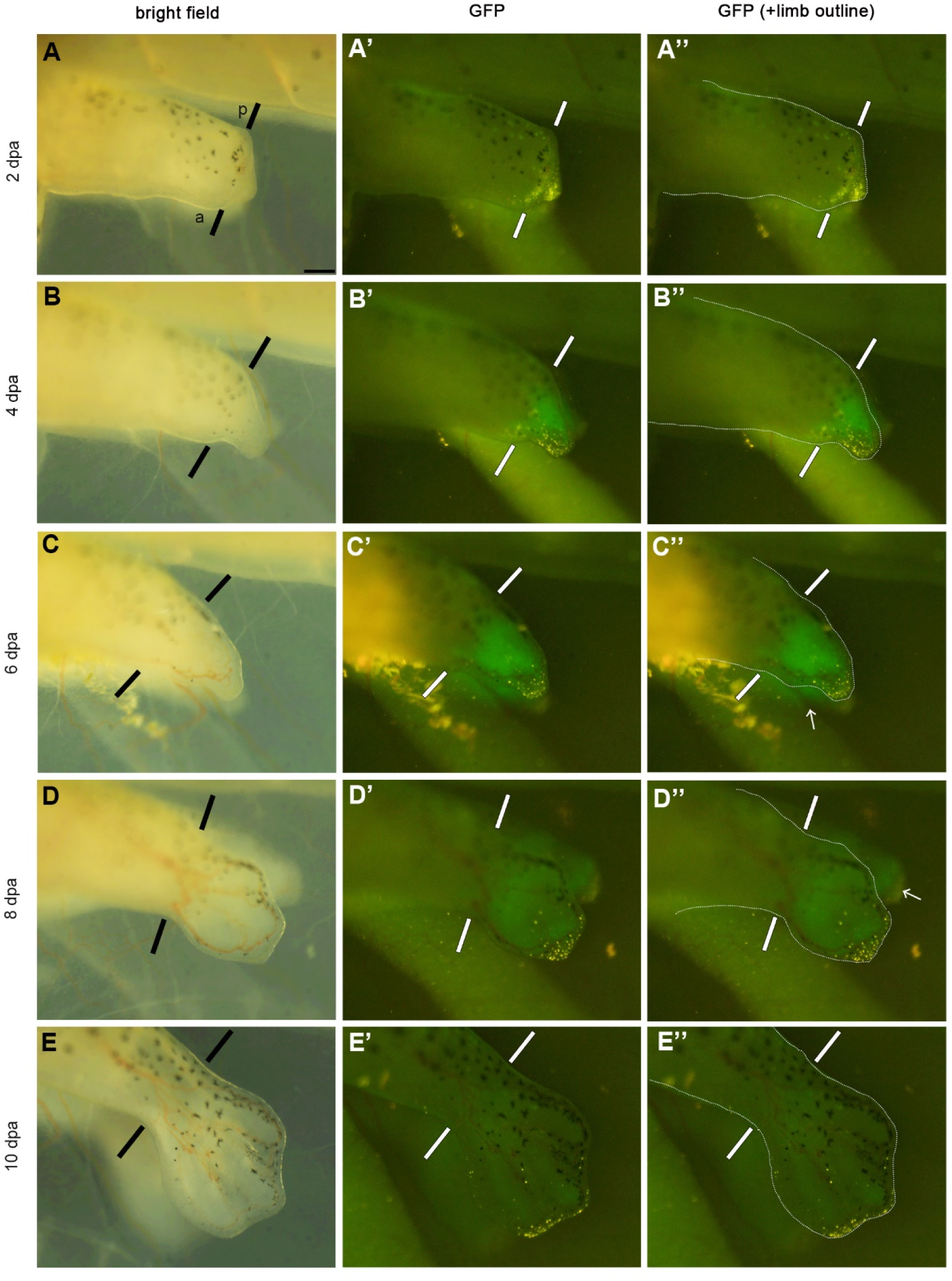
Spatio-temporal expression pattern of MFCS1-GFP in a regenerating limb of a young tadpole. All of the pictures are focused on a limb blastema of the same tadpole (identical to Fig. 3Q-T) of line 1. Left and right limb buds were amputated at the presumptive ankle level of a stage 54 tadpole. GFP fluorescence was distributed throughout the regenerating limb blastema at 4 dpa and 6 dpa. (A’’, B’’, C’’, D’’, E’’’) Limb blastemas and stumps are outlined by dotted lines. n=16 of 16 blastemas from a total of ten MFCS1-GFP tadpoles of line 1 and 6 of 6 blastemas from a total of three MFCS1-GFP tadpoles of line 2, respectively. A pair of lines, the amputation level: Arrows, GFP fluorescence of the contralateral right limb blastemas: a, anterior: p, posterior. Bar = 200 μm.

**Fig. S3.**
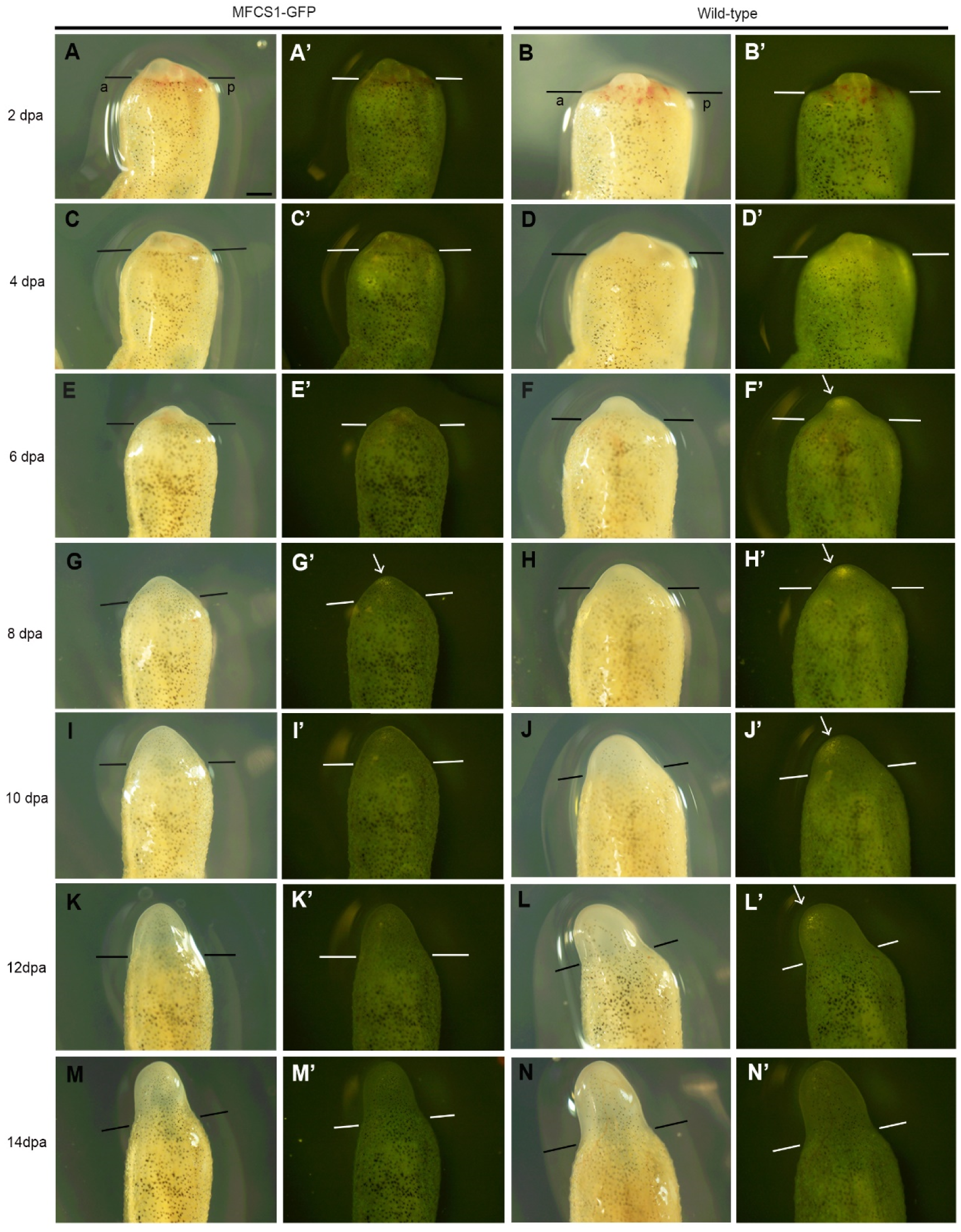
MFCS1-GFP expression is inactive in a limb blastema of a froglet. A blastema of the same froglet of line 1 (left two columns; identical to Fig. 3M-N) and a blastema of the same wild-type froglet (right two columns) from 2 dpa to 14 dpa. No specific GFP fluorescence was observed in the blastema of the Tg froglet at any timepoint. n=4 of 4 blastemas from a total of four froglets of line 1, 6 of 6 blastemas from a total of four froglets of line 2 and 8 of 8 blastemas from a total of six wild-type froglets as controls, respectively. A pair of lines, the amputation level: An arrow, non-specific fluorescence commonly observed both in the Tg froglet and wild-type one: a, anterior: p, posterior. Bar = 500 μm.

**Fig. S4.**
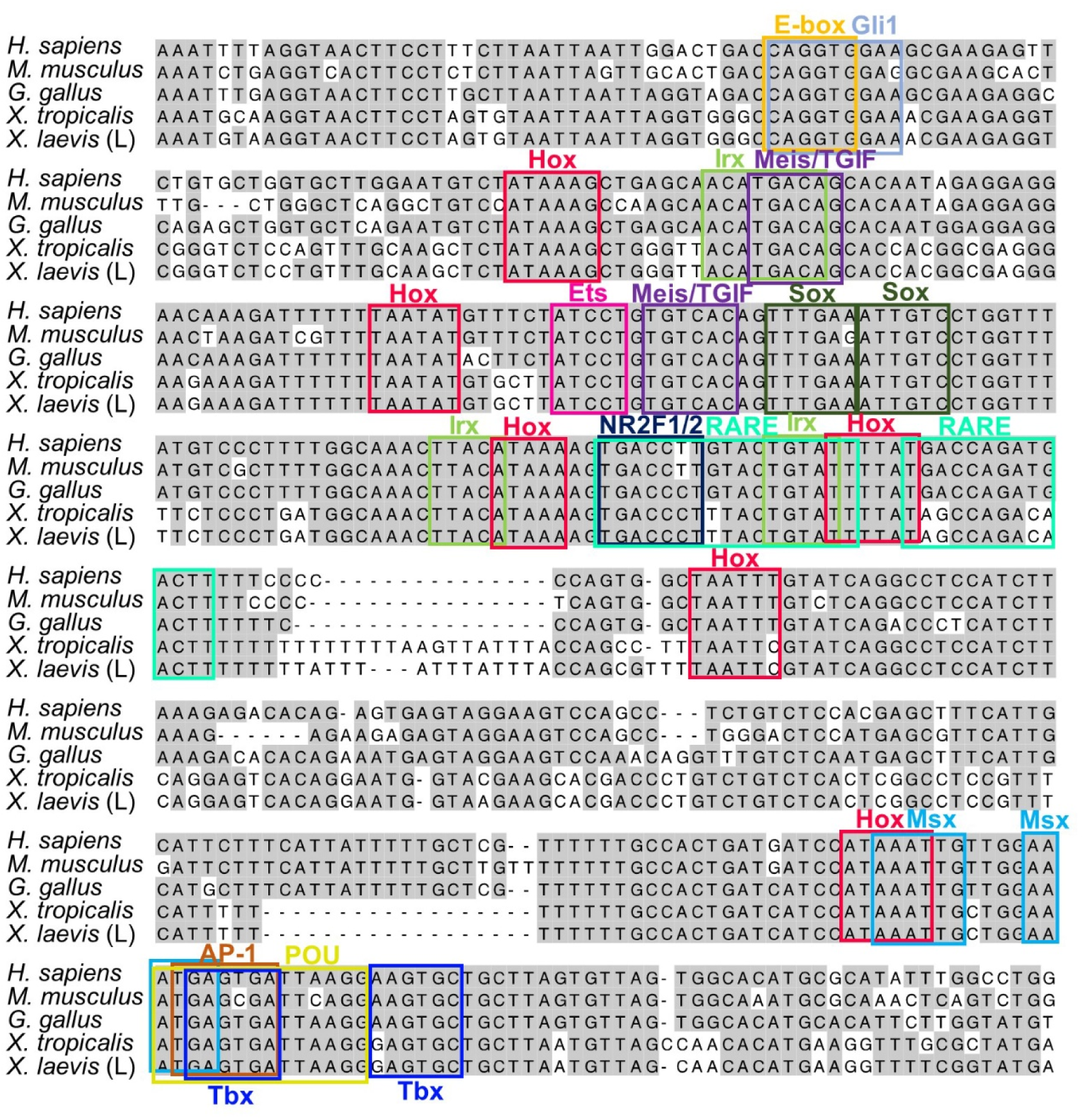
Alignment of MFCS1 highly conserved sequences (identical sequences shown in Fig. 4A) from the human, mouse, chick, *X. tropicalis* and *X. laevis*. Nucleotides identical in more than two sequences are shaded in grey. Conserved transcription factor-binding motifs (Hox and other transcription factors) are boxed.

**Fig. S5.**
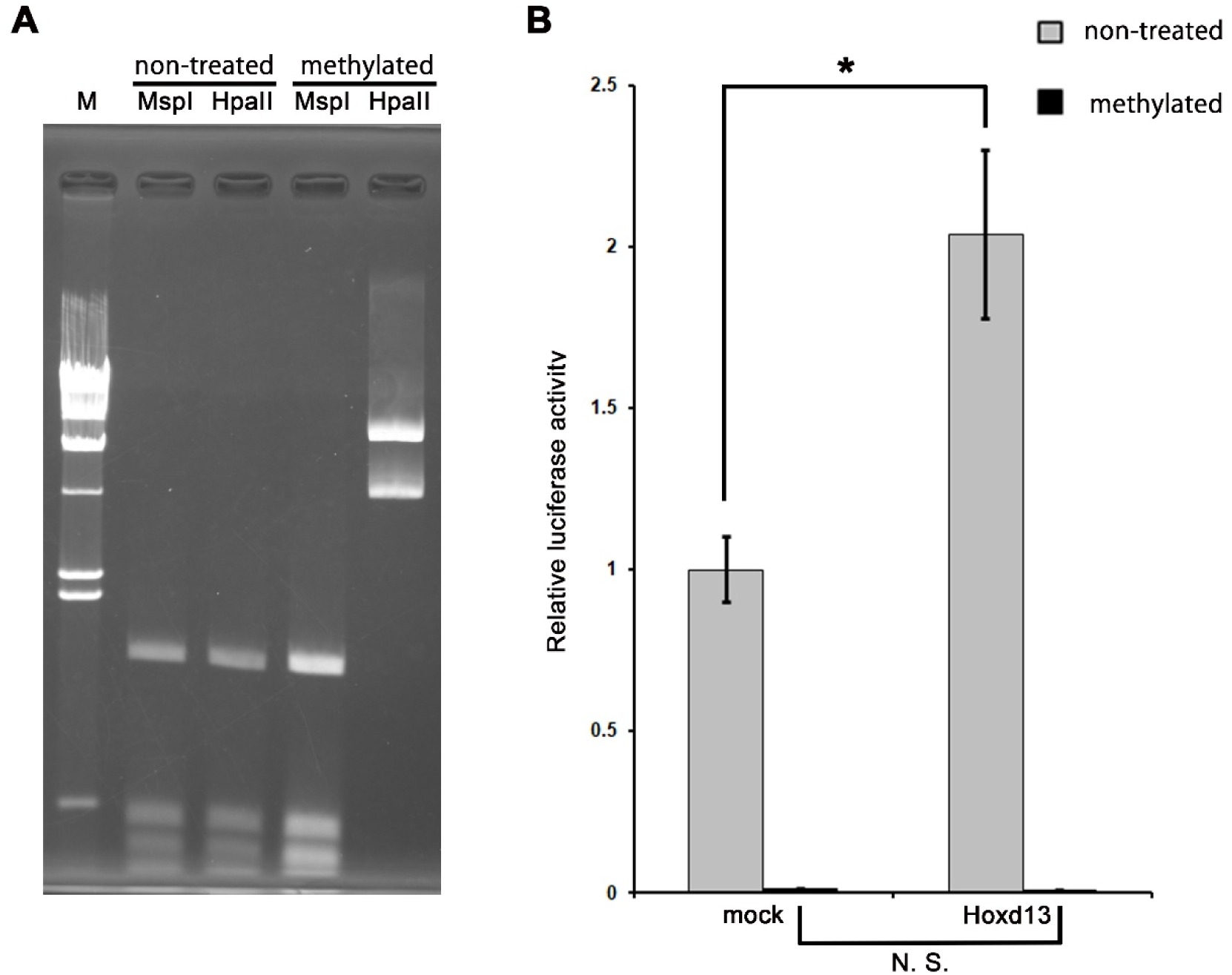
DNA methylation suppresses the response of MFCS1 to Hoxd13 *in vitro*. (A) Methylation-sensitive restriction digestion visualized by agarose gel electrophoresis. Plasmid DNA of the MFCS1-*shh* promoter-*luc2* reporter was treated with CpG methyltransferase (M. SssI, NEB) according to the manufacturer’s protocol. Methylation efficiency of the plasmid DNA was evaluated by digestion with methylation-sensitive (HpaII) and -insensitive (MspI) restriction enzymes. The CpG methylation prohibited the digestion of the plasmid DNA specifically with HpaII. M, λ/HindIII marker. (B) Luciferase activity in NIH3T3 cells transfected with the methylated or non-treated MFCS1-*luc2* reporter constructs and the *Hoxd13* expression construct. The mean value of luciferase activity in cells transfected with the non-treated reporter and mock expression constructs was set to 1.0. Error bars: standard deviation from the mean (n=3). While a significant increase of *Hoxd13*-induced luciferase activity was observed in cells transfected with the unmethylated reporter construct (*p<0.01), being consistent with Fig. 4B, there was no significant increase in cells transfected with the methylated reporter construct despite *Hoxd13* expression. N. S., not significant.

**Fig. S6.**
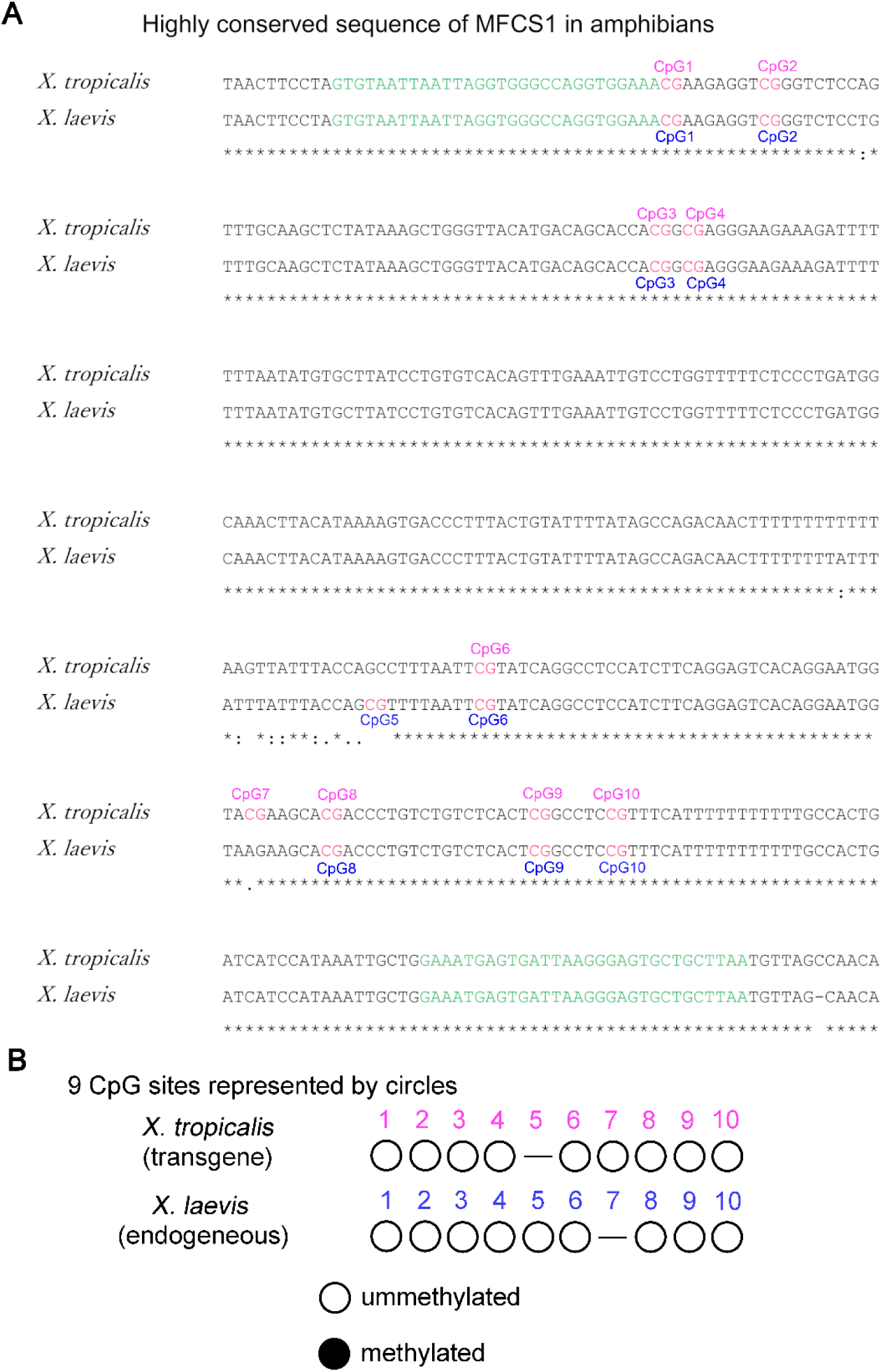
*X. tropicalis* and *X. laevis* have 9 CpG sites in the highly conserved sequence of MFCS1. (A) Alignment of MFCS1 highly conserved sequences between *X. tropicalis* and *X. laevis*. Green letters correspond to MFCS1_BS_Fw and MFCS1_BS_Rv primers for bisulfite sequencing (see also Table S3). (B) Open and black circles represent unmethylated and methylated CpG sites, respectively.

**Fig. S7.**
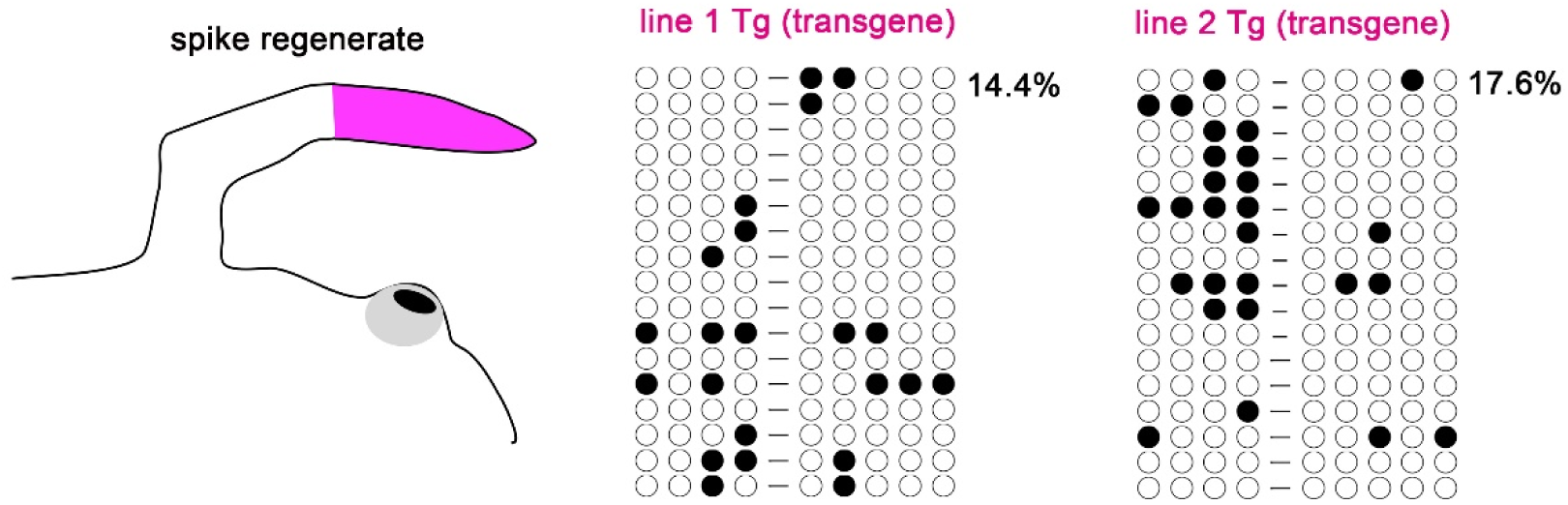
Methylation pattern of MFCS1 in a spike regenerate after limb regeneration of a froglet. After limb blastemas of froglets completed spike regeneration, regenerate samples (pink) were obtained from line 1 and line 2 Tg frogs.

**Table S1.**
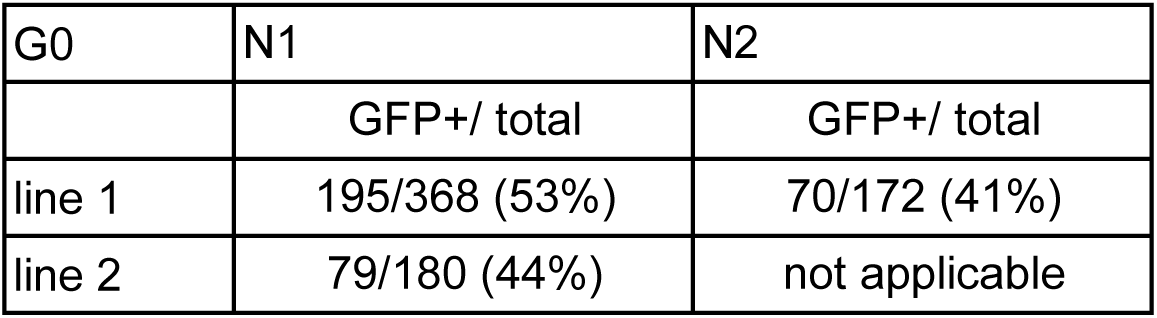
Rate of germline transmission and activity of MFCS1-GFP

**Table S2.** Supernumerary limb formation and ectopic GFP in rotated limb buds

| Sample | No. of original digits | No. of supernumerary digits | No. of digits (total) | Is there ectopic GFP? | Figure |
| --- | --- | --- | --- | --- | --- |
| 1 | 5 | 2 | 7 | Yes | - |
| 2 | 5 | 2 | 7 | Yes | - |
| 3 | 5 | 2 | 7 | Yes | - |
| 4 | 5 | 2 | 7 | Yes | - |
| 5 | 5 | 4 | 9 | Yes | Fig. 2F, G |
| 6 | 5 | 2 | 7 | Yes | - |
| 7 | 5 | 1 | 6 | Yes | Fig. 2D, E |
| 8 | 5 | 0 | 5 | No | - |
| 9 | 5 | 0 | 5 | No | - |
| 10 | 5 | 0 | 5 | No | - |
| 11 | 5 | 2 | 7 | Yes | - |

0° rotated limb buds (controls)
| Sample | No. of original digits | No. of supernumerary digits | No. of digits (total) | Is there ectopic GFP? | Figure |
| --- | --- | --- | --- | --- | --- |
| 1 | 5 | 0 | 5 | No | Fig. 2B, C |
| 2 | 5 | 0 | 5 | No | - |
| 3 | 5 | 0 | 5 | No | - |
| 4 | 5 | 0 | 5 | No | - |
| 5 | 5 | 0 | 5 | No | - |
| 6 | 5 | 0 | 5 | No | - |
| 7 | 5 | 0 | 5 | No | - |
| 8 | 5 | 0 | 5 | No | - |
| 9 | 5 | 0 | 5 | No | - |
| 10 | 5 | 0 | 5 | No | - |
| 11 | 5 | 5 | 10 | Yes | - |
| 12 | 5 | 3 | 8 | Yes | - |

**Table S3.**
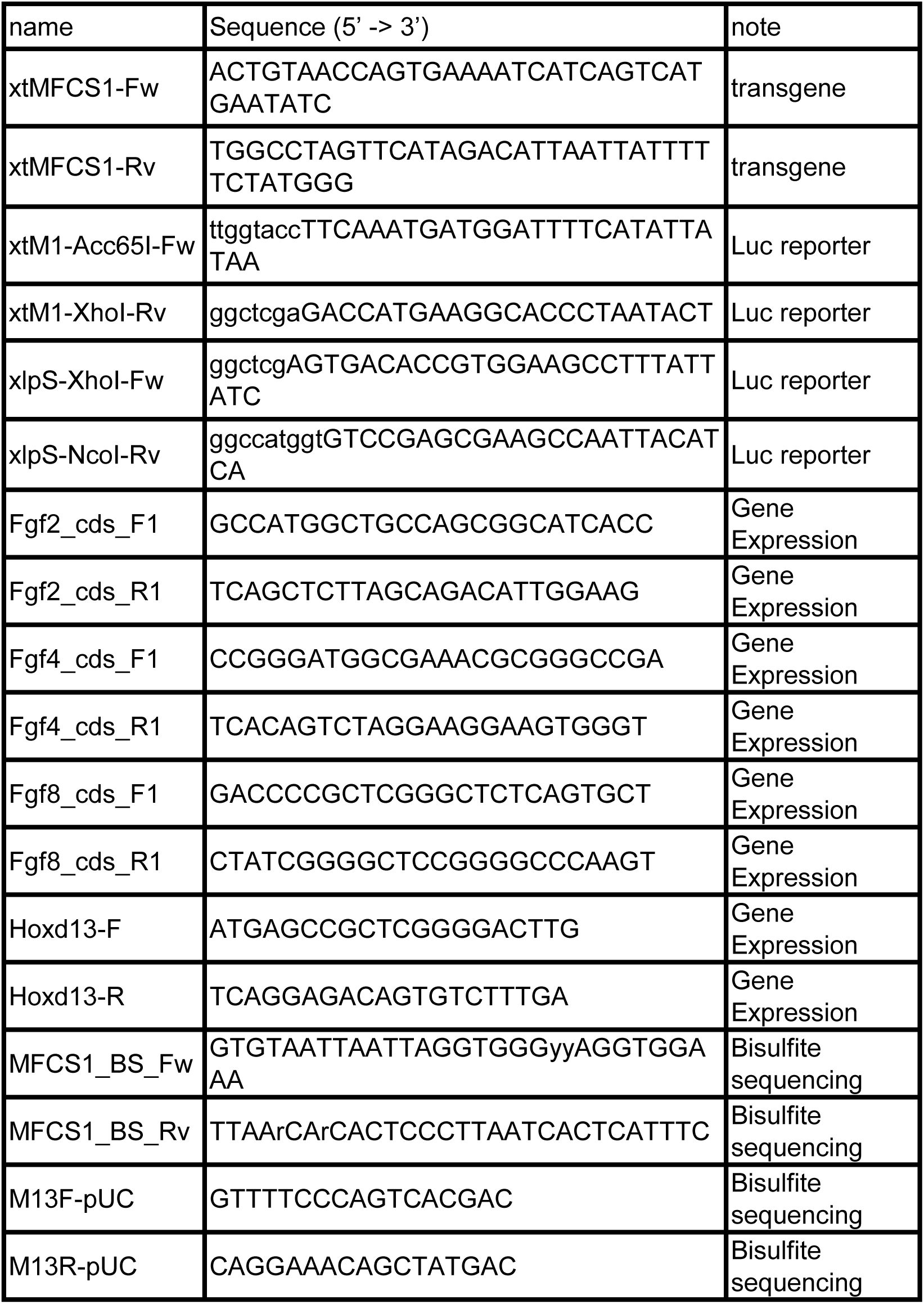
Primers for genomic- and RT-PCR

